# How do selective serotonin reuptake inhibitors effect metabolism? A lesson for teaching non-majors about enzymes and metabolic pathways

**DOI:** 10.1101/2025.11.20.689583

**Authors:** Trevor T. Tuma, Abigail M. Johnson, Peggy Brickman

**Affiliations:** Department Biochemistry & Molecular Biology, University of Georgia; Division of Biological Sciences, University of Georgia; Department of Plant Biology, University of Georgia

## Abstract

Understanding foundational concepts in molecular biology and biochemistry can be challenging for many college students. For non-science majors, an introductory biology course may be their sole exposure to college-level science and only opportunity to learn about enzymes and metabolism. Framing these topics within issues that are personally relevant can make the material more meaningful and impactful for non-majors. There is a need to develop curricula that instructors of large-enrollment introductory biology courses can use to teach non-biology majors about enzymes and metabolism. Here, we designed and implemented a lesson that integrates enzyme function and metabolism within the context of selective serotonin reuptake inhibitors (SSRIs). SSRIs are a widely prescribed class of antidepressants whose efficacy can be influenced by enzymes. Genetic polymorphisms in the genes encoding these enzymes can result in individual differences in metabolic responses and lead to medication-induced weight gain. Framing the content in this way allows non-major students to connect enzyme function to genetics, energy balance, and metabolic processes through a health-related, societally relevant issue. In our lesson, students engaged in collaborative learning activities that emphasized interpreting biological data, graphing, and scientific reasoning. Pre- and post-assessment data from a conceptual inventory showed learning gains that exceeded baseline data and were greater than those observed with the traditional curriculum. Students also reported a more coherent understanding of enzyme function and a greater appreciation of the content’s relevance to their own lives. This lesson provides an example of embedding biochemical concepts in real-world contexts for students who are non-science majors.

## INTRODUCTION

As part of their general education requirements, most undergraduates in the United States take at least one natural science course, which is often an introductory biology class^1^. For many non-science majors, this course may be their only exposure to college-level science. Many biology faculty find it challenging to engage this large population of students. Non-major students often differ from majors and are less likely to see science as relevant to their careers^2^, report lower motivation to learn science^3^, are less likely to view themselves as a science person^4^, and more prone to hold misconceptions about the nature of science^5^. Despite these challenges, non-majors will become future voters, educators, professionals, and caregivers, shape public discourse, and engage in decision-making on science-related issues. To better serve these students and foster a more scientifically literate public, there is a critical need to make non-major biology course content meaningful, relevant, and accessible for these students^6^.

One promising approach for overcoming these challenges is to frame biology content around issues that are personally and socially relevant to students. Evidence suggests that students are more likely to find science meaningful when they see its relevance to their communities, health, and well-being, and when they are afforded opportunities to engage with science beyond the classroom^6,7^. Students themselves have expressed a desire for more topic-based and concept-oriented courses, particularly within non-majors’ biology^8^. However, an analysis of 78 non-majors’ syllabi revealed that nearly half focus solely on biological content with little to no connection to societally relevant and socio-scientific issues^9^. Such a dominant focus on scientific content may limit non science major students’ ability to connect what they are learning to real-world problems and apply scientific thinking beyond the classroom.

One way to connect core biological content with real-world relevance relates to the teaching of metabolism and enzyme function, which are foundational for understanding core biological principles. Enzymes are fundamental biological catalysts that drive the majority of chemical reactions within living organisms. Understanding enzyme function and regulation is essential for broadening students’ understanding of the dynamic nature of metabolism, the mechanism(s) underlying physiological processes, and the maintenance of homeostasis. These concepts are also central for understanding topics in biotechnology, precision medicine, and pharmacology. Furthermore, the importance of metabolic pathways and their regulation is reflected in the millions of publications that investigate these biological processes^10–12^. Ongoing rapid advancements in metabolic engineering have further emphasized the role of enzymes in synthesizing fuels, pharmaceuticals, and other valuable biproducts^13^. More practically, understanding how enzymes function is essential for understanding how biochemical pathways can impact human health and disease.

Despite their centrality, understanding concepts in molecular biology and biochemistry are often challenging for both major and non-major students^14–16^. These topics require abstract reasoning and systems thinking. Often, they also involve concepts such as free energy, feedback inhibition, and biochemical interactions. While biochemistry students typically have opportunities to develop this knowledge across multiple courses and can integrate multiple ideas in ways that allow them to leverage this knowledge to understand metabolism^17^, non-majors may encounter them for the first and only time in a single semester. Non-majors may also benefit from pedagogical approaches that make learning science learning useful and relevant, rather than prioritizing content breadth or technical depth.

Efforts to incorporate the teaching of enzymes and metabolism within real-world contexts has the potential to improve non-majors learning experiences. One way to contextualize enzyme function and metabolism within a relevant societal issue relates to the use of selective serotonin reuptake inhibitors (SSRIs). SSRIs are a class of widely prescribed medications used to treat many mental health conditions, including depression, anxiety, obsessive-compulsive disorder, and post-traumatic stress disorder (PTSD)^18^. In the United States, more than one in ten adults takes an anti-depressant, and SSRIs are the most commonly prescribed type of antidepressant^19^. Their use has increased since the COVID-19 pandemic, particularly among young adults^20^.

SSRIs function by inhibiting the serotonin transporter protein (SERT), which is responsible for serotonin reuptake into the presynaptic neuron after it has been released in the synaptic cleft. Normally, after serotonin is released and transmits its signal to a neighboring neuron, the serotonin transporter protein brings it back to the original presynaptic neuron. By blocking this transporter protein, SSRIs increase serotonin levels in the synaptic cleft which enhances neurotransmission and, in many cases, alleviates symptoms of depression and anxiety.

Although SSRIs directly target transport proteins, their efficacy is heavily influenced by metabolic enzymes. Many SSRIs are metabolized by liver enzymes belonging to the cytochrome P450 enzyme family, such as CYP2D6 and CYP3A4. The same enzymes also play roles in the breakdown of neurotransmitters. In the brain, excess serotonin not reabsorbed by SERT can be broken down by monoamine oxidase (MAO). MAO is an enzyme that metabolizes serotonin into inactive compounds and acts as an example of how enzymes can regulate levels of a neurotransmitter.

Genetic variation in the genes that encode these enzymes (e.g., polymorphisms in CYP2D6 or CYP3A) can also influence how efficiently an individual metabolizes SSRIs. These genetic variations can lead to slower or faster drug metabolism, which alters how much of the medication stays in the body and for how long. For example, individuals with non-functional CYP2D6 alleles may metabolize SSRIs slower, resulting in prolonged drug exposure and increased side effects, such as weigh gain. Such genetic variability provides another opportunity for students to learn about the role of metabolism in shaping individual responses to antidepressant treatment. In addition, the connection between enzymes, metabolism, and genetics highlights the role of metabolic regulation within the context of real-life health decisions.

In this manuscript, we present a multi-class teaching lesson designed to integrate enzyme function and metabolism within the context of SSRIs. Our goal in developing this lesson is to support college non-major biology students in developing foundational content knowledge and the scientific literacy needed to engage with biochemical and metabolic topics beyond the classroom. By leveraging a topic that is timely, personal, and socially relevant, this lesson aims to deepen non-major students’ conceptual understanding. Our approach aligns with recommendations from Vision and Change and American Society for Biochemistry & Molecular Biology educational frameworks that emphasize core biological concepts, such as “pathway transformations of energy and matter”, “energy and metabolism” and “systems thinking”^8,21^.

### Intended Audience

Students of all years at the college level who are taking a non-majors biology course.

### Required Learning Time

We developed this lesson to cover two 50-minute class periods. Students complete pre-class homework assignments (Supporting Files S1 & S2) that review concepts related to enzyme structure and function prior to the first lesson and enzyme regulation prior to the second lesson. Students complete additional worksheets (Supporting Files S3 & S4) in class as they work through the lesson. Optional activities intended to generate more in-depth exploration and discussion can easily be added.

### Prerequisite Student Knowledge

Students should have a basic understanding of the structure and function of biological molecules, including the ability to identify and describe proteins. Students also benefit from the pre-class background reading on enzymes.

### Prerequisite Teacher Knowledge

Before teaching this lesson, instructors should feel confident in their understanding of basic enzyme concepts, including substrates, products, and enzyme function in metabolism. They should also be familiar with human metabolic pathways and particularly those involved in medication metabolism. Instructors will also benefit from understanding the pharmacology of SSRIs, including their mechanism of action in the brain, potential metabolic side effects, and how changes in enzyme activity can influence drug efficacy and physiological outcomes. To support students’ ability in analyzing figures and graphs, instructors might explore a graphing lesson that engages students in stepwise, scaffolded activities with explicit modeling, hand-sketching, and collaborative interpretation of data^22–24^.

## SCIENTIFIC TEACHING THEMES

### Active Learning

In this lesson, students are engaged in multiple forms of active learning. Before class, students complete assigned readings to introduce key content and foundational concepts. These pre-class readings are intentionally designed to reinforce content knowledge and prepare students for in-class collaborative work.

During class, students work in groups of four to complete worksheets that ask them to interpret graphical representations of real biological data and make predictions based on their analyses^25^. Nearly every component of the in-class assignment involves discussion and collaboration, which prompts students to articulate their thinking^26^. Peer interactions through group work have also been shown to promote deeper engagement with the content^27^. The group assignments are purposefully structured to foster task interdependence and individual accountability^25^. For example, students work collaboratively to complete worksheet questions and identify an SSRI for their final project. Each group member is assigned specific responsibilities and tasks, which helps ensure that all students in the group contribute meaningfully to the tasks and mitigates social loafing. As part of the project, students are asked to describe their individual contribution which further reinforces accountability.

Throughout the lesson, students are given multiple opportunities to write, verbalize, and articulate their thinking about biology throughout the lesson^28^. One way this is integrated is through think-pair-shares with groupmates. For example, the instructor poses a question and allows students time to think individually about how they would answer the question. Students are then asked to discuss their reasoning with their group. Then they share their answer by submitting a response to a clicker question. The use of clicker questions provides instructors with real time evidence of all students’ thinking, and particular response choices can serve as the basis for class discussions to clarify student misconceptions^29^. Students are instructed to explain in their groups not only why the correct answer is right, but also why the other options are incorrect. These prompts encourage reasoning, which helps students connect their thinking to evidence^30^. When a clicker question reveals a range of answers, the instructor can facilitate a debrief discussion to clarify misconceptions before students’ reattempt answering the question.

Students are provided with opportunities to develop their metacognitive skills, or their ability to monitor and regulate their own learning. Students are given strategies to help them become more metacognitive. For example, students are given learning objectives and are encouraged to answer the objectives as self-assessment questions after the lesson is over and as they study for summative assessments^31^. Students are also provided with practice exam questions to help study and identify what concepts they understand and do not understand. Students can also access their responses to the clicker questions after class, which they can use to guide their subsequent studying.

Through social metacognition in the lesson, or the awareness and regulation of others’ thinking^32^, students can monitor both their own understanding and that of their peers. Students can verbally correct their own thinking and the thinking of their groupmates during group discussions. For example, they can evaluate the explanations offered by their peers and identify areas where their thinking is in alignment or potentially divergent. By doing so, students can deepen their own conceptual understanding while contributing to the groups’ collective learning.

### Assessment

This lesson includes multiple opportunities for formative and summative assessments. Both are mapped to unit learning objectives.

### Formative Assessment

Students’ understanding of enzymes was assessed using a concept inventory administered pre-and post-lesson to measure learning gains (Supporting File S5). Prior to class, students completed a reading assignment and answered a series of multiple-choice questions based on this reading to prepare for in-class activities. During class, they worked collaboratively on worksheets that expanded on the learning objectives.

On the final day of class, students complete a group assignment exploring enzyme-mediated metabolism of antidepressant medications (Supplemental File S6). Each group is tasked with selecting one antidepressant from a provided list. Groups are tasked with identifying enzyme-substrate-product relationships, considering the factors that could alter enzyme activity, and evaluating the impact on drug efficacy and dosing. They also constructed graphs representing how different metabolizer types (e.g., poor, normal, ultrarapid) would affect enzyme activity and explained how these differences could affect drug treatment. Students were also asked to support their answers with at least two high quality, peer-reviewed sources. Students also completed reflective assessments designed to encourage metacognition and personal connection to the content. Students responded to open-ended questions asking them to identify challenging and confusing aspects of the lesson and to consider the factors contributing to these difficulties. They also reflected on how the concepts they learned about enzymes and metabolism connected to their prior knowledge, as well as any potential applications of the content to other courses and real-world contexts.

### Summative Assessment

Summative assessment includes 4-5 multiple choice questions administered on a cumulative end of unit exam (Supporting File 7).

### Inclusive Teaching

The lesson begins with a hook designed to capture students’ interest in the content and establish the relevance of enzymes and metabolism. Students are first shown data illustrating the prevalence of depression across the United States. The data reveals that the prevalence of depression has increased over time, particularly during the COVID-19 pandemic. This opening grounds the topic in a real-world context and acknowledges that depression is a widespread public health issue.

Students are then introduced to data showing the relationship between SSRI use and one of their documented side effects, weight gain. Presenting this connection encourages students to think about the complexity of biology and human health while also recognizing that individuals can have varied physiological responses to medications. Relating biology concepts, particularly enzymes and metabolism, to real world issues allows students to see the relevance and purpose of the content to their own lives, which can promote learning and persistence^33^.

Integrating relevance into biology content is thought to be particularly impactful for students from traditionally marginalized and underrepresented backgrounds ^34,35^, and thereby can lead to more equitable learning outcomes.

To further promote an inclusive educational environment, students work collaboratively in small groups on assignments. Group work can foster a sense of community by encouraging students to discuss their thinking, share their perspectives, and compare their reasoning processes. Importantly, students work in groups that are assigned randomly rather than self-selected, because instructor-assigned groups can improve learning outcomes and help to ensure that all students are included^36,37^. This approach also helps present instances where some students are excluded from groups and therefore supports a more inclusive classroom environment^38^.

We also recognize that conversations relating to depression and mental health can be emotionally challenging, and students may have varied prior exposure to such conversations. At the beginning of the lesson, we include a content warning and an acknowledgment that some of the material may be sensitive for some students. Students are encouraged to communicate with the instructor if the content becomes too difficult to engage with and, if necessary, may request an alternative assignment.

## LESSON PLAN

### Student Preparation Before Day 1

Student preparation for this lesson can include two components: a brief pre-lesson assessment and a short pre-class reading. An optional short (taking ∼10 minutes) multiple-choice conceptual inventory can be administered either online or in person to determine student knowledge (Supporting File S5). Second, students should read open-source textbook content on metabolic pathways and enzymes. The reading introduces the concept of metabolism as a series of energy-transforming reactions within cells and explains the roles of anabolic and catabolic pathways in energy balance. The section concludes with an overview of enzyme structure and function and explains how enzymes can act as catalysts. We included four self-assessment items following the reading to help students check their comprehension and apply these concepts prior to class (Supporting File S1).

### In-Class Day 1

Day 1 introduces students to the role of neurotransmitters in mood regulation through a brief lecture (Supporting File S8). The lecture also connects these ideas to the clinical relevance of depression. The instructor presents slides that illustrate how serotonin is synthesized in the body. The instructor introduces the ideas of metabolic pathways and medication interactions by describing the enzyme mediated conversion of tryptophan to 5-hydroxytrptophan by tryptophan hydroxylase. Students begin by answering clicker questions drawn from the worksheet (Supporting File S3). The first question prompts students to identify the substrate, enzyme, and product in the serotonin synthesis pathway.

Students are then prompted to think about how substrates, enzymes, and products may change over time as the reaction progresses by interpreting data from a graph that depicts concentrations of the substrate and product over time. Students are then asked to fill out a table predicting how these concentrations of tryptophan, 5-hydroxytrptophan, and tryptophan hydroxylase will change over time based on this information. This question prompts students to make quantitative predictions of concentrations based on their understanding of enzyme function. Students then complete another question asking them to interpret the role of tryptophan hydroxylase in this conversion. These activities are designed to explain the roles and dynamic interplay of enzymes, substrates, and products.

The instructor then presents slides with data highlighting that depression remains a major health concern and that only about 65% of patients respond to antidepressants. Students are prompted to think about why medication effectiveness may vary. This leads naturally into content examining the biological processes that occur when SSRIs are ingested. Students are then asked to draw a diagram tracing what happens to an antidepressant from ingestion to elimination. They are asked to include absorption, distribution, metabolism, and excretion in the drawing. This activity helps them visualize how the body processes medications and how this may be related to metabolic pathways.

After students complete their diagrams, the instructor explains what happens biologically when antidepressants are consumed by using fluoxetine (Prozac) as an example. The instructor describes how fluoxetine is metabolized in the liver by the enzyme CYP2D6 into active metabolites, such as norfluoxetine, before eventually being excreted through the kidneys.

Students are then asked to make predictions about why some individuals may respond differently to antidepressants. Students are encouraged to connect their reasoning to how the body breaks down and processes medications and how this could influence medication effectiveness. The next question on the worksheet asks students to examine a graph showing antidepressant concentration over time. Students first interpret what the graph is showing by identifying the x-axis and y-axis before describing the trends they see in the data. They are asked to consider what might cause the concentration to rise initially and why it eventually decreases. This question encourages students to connect their understanding of absorption, metabolism, and excretion to changes in antidepressant levels.

The instructor then introduces enzyme inhibition through a short lecture on how antidepressant medication can interfere with metabolism. Using sertraline as an example of the metabolic breakdown of SSRIs, the instructor explains how different liver enzymes can act in parallel or in series to catalyze its breakdown into demethylsertraline, which is ultimately excreted by the kidneys. The lecture then introduces Omeprazole (Prilosec), a proton pump inhibitor commonly used to treat acid reflux. Omeprazole is known to inhibit tryptophan hydroxylase and thereby potentially interfere with SSRI metabolism. To illustrate the mechanism of inhibition, the instructor shows a visual comparison of normal and inhibited enzyme activity. One molecular visualization shows a typical enzyme reaction in which the enzyme binds to a substrate and releases a product. The second visualization shows how the presence of an inhibitor blocks the substrate from binding to the enzyme’s active site. The instructor also explains that Omeprazole remains on the market because it inhibits MAO, the enzyme that normally breaks down serotonin, which can counteract some of the side effects on serotonin metabolism.

Following this explanation, students examine data showing concentrations of tryptophan, 5-hydroxytrptophan, and tryptophan hydroxylase over time in the presence or absence of Omeprazole. Students use the data to interpret what happens to 5-hydroxytryptophan levels when Omeprazole is present and what can be concluded about Omeprazole’s role in the enzyme pathway for serotonin metabolism. These last questions assess student understanding of the consequences of enzyme inhibition on substrate and product formation. The class concludes by debriefing the questions and an explanation of how inhibition disrupts metabolic pathways and can alter SSRI effectiveness.

### Student Preparation Before Day 2

Before the second day of the lesson, students read open-source textbook material on enzyme inhibition to build on their prior understanding of enzyme structure and function (Supporting File S2). The reading provides an introduction to the mechanisms of enzyme regulation, including competitive and noncompetitive inhibition. It also connects these biochemical processes to the development of pharmaceutical drugs that target specific enzymes. The reading concludes with an overview of feedback inhibition and the role of cofactors and coenzymes in regulating metabolic pathways. To assess comprehension, students answer two questions that ask them to identify the effect of enzyme inhibitors on enzyme activity and to apply these concepts to examples of drug-enzyme interactions (Supporting File S2).

### In-Class Day 2

The second-class session begins with the instructor reviewing the concepts from the previous lesson (Supporting File S9). They instruct students that today’s class will focus on connecting enzyme inhibition and antidepressant metabolism to the physiological effects of SSRIs. The instructor introduces results from data showing that weight gain is a very common side effect among antidepressant users. The lecture slide shows data depicting medication-induced weight change across several common SSRI treatments. The instructor leads a short discussion on how these weight-related outcomes may be linked to serotonin metabolism and cues students to think about how enzymes may be associated with this process.

The instructor then explains that SSRIs increase serotonin levels in the brain. Increased serotonin levels can positively enhance mood but also influence other enzymes that regulate appetite and energy storage. Students are then asked to complete a multiple-choice question that asks how serotine levels might influence enzyme activity related to appetite regulation and metabolism (Supporting File S4). The instructor then reviews the options with class and explains that increased serotonin can alter enzyme activity in ways that affect hunger and metabolic rate, thereby potentially contributing to weight gain in some individuals.

The instructor then transitions to the topic of genetic variability in medication metabolism. The class is provided with examples of genetic testing for SSRI metabolism as a tool for identifying differences in how individuals may process antidepressant medications. The instructor explains that individuals can be classified as ultrarapid, normal, intermediate, or poor metabolizer, which can influence both the effectiveness of SSRIs and the likelihood of experiencing side effects such as weight gain.

To illustrate this concept, students are shown biology data illustrating how variations in genes encoding cytochrome P450 (CYP) enzymes can be associated with different metabolizer types. The graph displays changes in body weight over time for individuals with different CYP2C19 metabolizer phenotypes. Students are then asked to interpret trends in the graph to understand how metabolic rate affects medication concentration and physiological responses. Students complete a section on their worksheet using data from the graph to summarize both the potential weight gain response and to draw conclusions about what this means for each metabolizer type. Through this question, students should understand that variation in the CYP enzyme can influence their metabolizer type and how efficiently SSRIs are broken down in the body. This, in turn, affects medication concentration and the likelihood of experiencing side effects such as weight gain.

Students then apply their understanding of how CYP enzyme activity may influence fluoxetine concentration. Specifically, they are asked to consider how enzyme activity would differ between a fast metabolizer and a slow metabolizer, and to predict the effects of these metabolizers on Sertraline (Zoloft) breakdown. Students are then asked to consider first the x- and y-axis for each graph and then depict how enzyme activity may vary over time between fast and slow metabolizers after consuming Sertraline. These questions are designed to help students visualize the relationship between enzyme activity, medication metabolism, and the resulting physiological effects.

The instructor continues with the Sertraline example by introducing CYP450 enzyme mutations to bring attention to their impact on SSRI metabolism. Students learn how variations in the CYP450 gene can affect the breakdown of Sertraline into its metabolite, desmethylsertraline, and influence drug levels in the body. Students then answer questions about the consequences of slower or less efficient enzyme activity, including how mutations in the gene that encodes the CYP450 gene could require adjustments in Fluoxetine dosage. Finally, students are also asked to consider how individuals who have a mutation in the CYP450 gene might make them a slower metabolizer of Sertraline. They are prompted to think about how this could affect drug levels in the body, the body’s response over time, and ultimately an individual’s weight gain.

### Day 2 Group Assignment

The remainder of Day 2 is dedicated to completing a group assignment that allows students to apply concepts from the in-class activities on SSRIs, serotonin metabolism, and enzyme variation (Supporting File S6). This group assignment has two purposes. First, it helps students apply principles of biology to support real-world decisions, such as reviewing potential adverse medication reactions. Second, it encourages students to explore general metabolic processes that the body uses to metabolize medications and to examine the roles that enzymes play in these processes.

This assignment helps students practice several key skills. First, the skills include identifying and evaluating the credibility of scientific sources, developing and clearly communicating scientific ideas in written form for a general audience, and planning, managing, and completing a project as a collaborative group. By integrating these skills into the group assignment, students gain practical experience that is relevant for real-world scientific skills. The lesson also reinforces foundational knowledge covered during the lesson. Specifically, students review the characteristics of enzymes that affect their function, learn to predict how changes in enzyme activity (e.g., inhibition, mutations) could influence levels of substrates and products, and explore different types of enzymatic regulation and inhibition.

After the lecture on SSRIs, weight gain, and metabolizer types earlier in the class, the instructor introduces the group assignment that will occupy the remainder of Day 2. The instructor provides a general overview of the assignment and that it will require students to identify and utilize high-quality sources of scientific information. The instructor provides an overview on how to use Google Scholar as a tool for identifying primary sources. The instructor also briefly explains the difference between different types of sources (e.g., primary, secondary) and how to evaluate their credibility.

Students then begin working on the group assignment (Supporting File S6). They continue to work in the groups that they were assigned during class and select one antidepressant medication from a provided list. Each group member is assigned to complete a different section of the assignment. Group member responsibilities are described below.

- Group Members 1 and 2 research, summarize, and cite sources for background information about the antidepressant their group selected (Questions #1-5 of the assignment).
- Group Member 3 explains the expected concentration of the antidepressant over time based on the scenarios that were covered in class (Questions #6-8 of the assignment). They also review the content from Group Members 1 and 2 for accuracy.
- Group Member 4 interviews all group members to gather responses to the last two questions, checks and edits the final responses to all questions, and submits the completed assignment.

Once students are oriented to the group assignment and the group roles are established, student groups investigate their selected antidepressant and work towards completing the assignment. The assignment asks them to explain how the drug is metabolized in the liver, to identify the main enzymes responsible for its breakdown, to determine the substrates and products of these enzymatic reactions, and to consider changes that could alter the enzyme metabolism of the antidepressant. Students are required to support their responses to these questions with at least two high-quality sources. They also are prompted to explain why they found the source to be relevant for the claim they are making.

The assignment also involves a data interpretation component. Students either create or analyze graphs showing how SSRI concentrations vary over time for different metabolizer types (e.g., underactive enzyme, ultrarapid metabolizer, or normal metabolizer). Students are asked to explain how variations in enzyme activity could result in a range of different physiological outcomes as well as result in potential side effects.

Finally, students complete several reflection questions describing what they found challenging, how what they learned relates to their existing knowledge, and how the content is useful and relevant for their lives; all of which are important for promoting student learning. Students are provided time to start the assignment during class, and any questions not completed during class are assigned as homework. Students document their contributions on the assignment to facilitate collaboration and accountability.

## TEACHING DISCUSSION

### General Observations

This lesson was implemented to multiple sections of a large enrollment (>180 students) introductory biology course for non-majors in January and September 2025. Overall, students appear to remain engaged and interested in this lesson. Many students have shared their personal experiences with antidepressants and weight gain with us. Other students have related the content of SSRIs to their coursework in their psychology and public health majors.

### Pre-Post Changes in Knowledge

In each section of the course, we assessed students learning about enzymes and metabolism (Supporting File S5). We assessed learning use items that were adapted from Gen Bio-Maps and American Society for Biochemistry & Molecular Biology questions to develop an eight-question, multiple-choice test^39,40^. The assessment was administered to students prior to the lesson (pretest) and at the end of the lesson (posttest). Students earned nominal course credit for completing the assessments. A total of 316 students across three sections of the course consented to use their data and completed both the pretest and posttest. We calculated each student’s average normalized learning gain from pretest to posttest^41^.

Students showed gains in their conceptual understanding of enzymes and metabolism (Figure 1). The mean learning gain for the two sections in which this lesson was implemented was 0.59 (SD = 0.38) and 0.56 (SD = 0.40). In comparison, mean learning gains were lower in the section taught with the traditional curriculum 0.43 (SD = 0.47).

**Figure 1.**
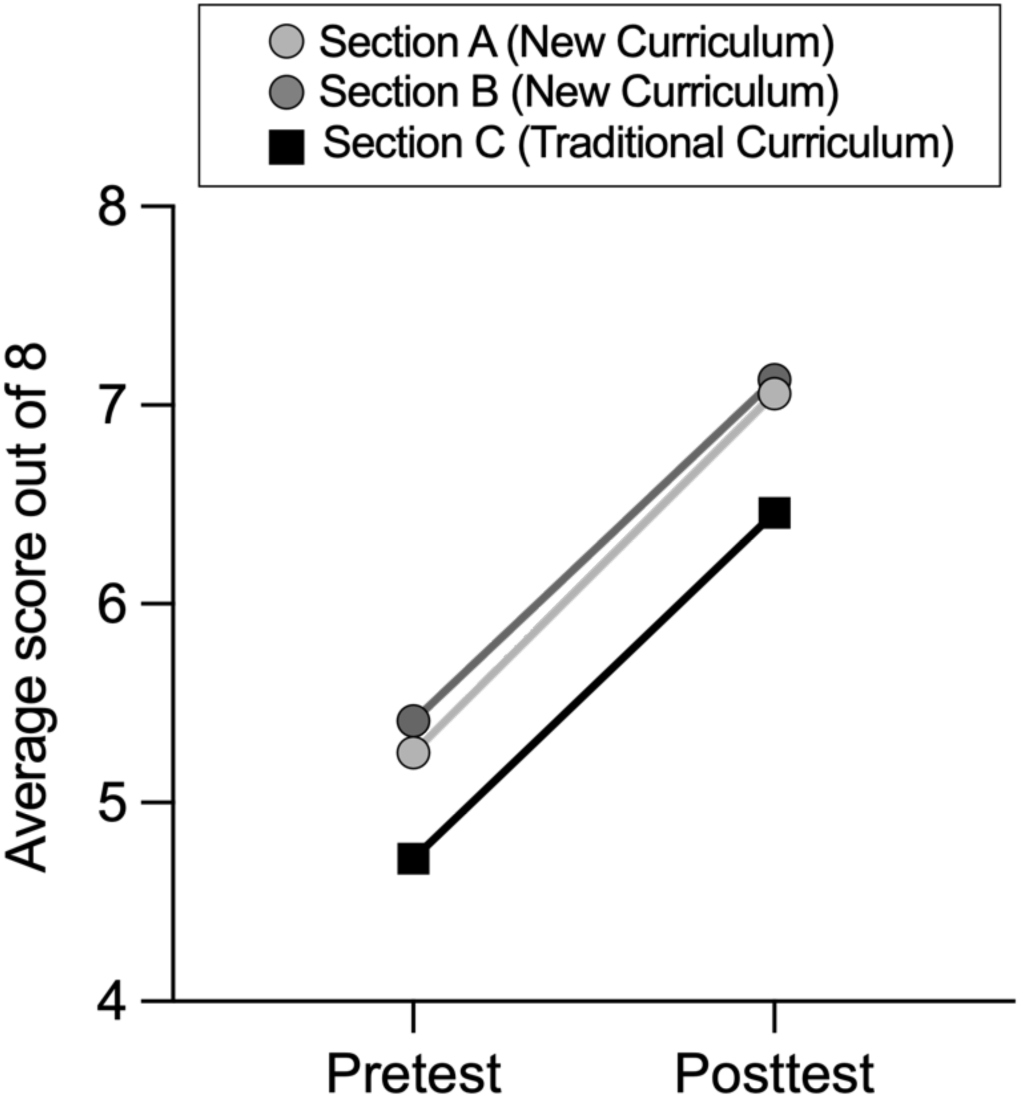
Students improved on the enzyme and metabolism assessment. Lines show the mean pretest and posttest scores (out of eight) on a conceptual inventory of enzymes and metabolism for each section. Two sections implemented the new lesson (Sections A & B; grey lines), while one section followed the traditional curriculum (Section C: black line). Student learning gains in sections taught with newly developed materials (black lines) compared to those in another section of the same course taught by a different instructor (grey line). Steeper lines indicate greater learning gains. The average normalized learning gain in sections with the newly developed materials was 0.59 and 0.56, while the average gain in the other section is 0.43.

### Student Responses to the Lesson

#### Challenges identified by students

To understand the challenges students reported during the lesson, we collected open-ended responses from students asking them to describe the part of the lesson they found most difficult and the factors that contributed to those difficulties. Student responses highlighted both conceptual difficulties and instructional factors that impacted students’ understanding. The prompt was also designed to encourage some metacognitive reflection, as students considered not only what was challenging, but also how and why these difficulties arose.

Responses highlight areas where students encountered difficulties, such as:

> *“One subject that took more time to understand was genetic variations and how polymorphisms in enzymes produce varying types of metabolizers. It was not a subject I have learned much about before, so it was a new concept to understand.”*
>
> *“One of the most challenging topics to understand is how SSRI inhibitors work in helping serotonin levels. To use, for example escitalopram, we found it confusing how it increases serotonin levels through inhibition (usually a word that is not correlated with increasing levels.) We later learned that it was because escitalopram blocks serotonin reuptake, that creates more serotonin in synaptic cleft so that more serotonin is built up. We feel that the idea of an ‘inhibitor’ was confusing as to how it helps build serotonin levels.”*
>
> *“I think the most challenging or confusing part of this unit was the topics covering SSRIs and medicinal drug metabolism. I think what contributed to making those topics difficult was the different possible metabolizers (slow, intermediate, extensive, and ultrarapid metabolizers), the different graphs (concentration vs. time, weight vs time, etc.), and the variety of SSRIs and the enzymes that metabolizes and breaks them down.”*
>
> *“The most challenging part was researching a new drug and comparing it to those discussed in class. It was difficult to connect the new drug’s substrates and enzymes to what was learned, especially given the complexity of their names. It was necessary to understand that, despite their names, these substances operate within the same categories of substrates and enzymes. Comparing them to real-world drugs like Lexapro also contributed to the difficulty.”*

#### Connections to prior knowledge on enzymes and metabolism

To understand how students integrated new content with their existing knowledge, we collected open-ended responses to a question asking students to explain how the material in the lesson connected to their existing knowledge about enzymes and metabolism. Research from cognitive and educational psychology emphasizes that learning is more likely to be sustained and transferable when students are prompted to connect new information to prior knowledge^42^. Encouraging these connections can help students organize complex concepts, such as enzyme regulation and metabolic pathways, into coherent mental models rather than treating them as isolated facts. Many students were able to link new information about enzyme function and metabolic pathways to concepts they had learned previously in other college or high school science courses. Other students noted that prior courses had introduced the terminology and provided a somewhat superficial description of the biological processes, but had not introduced deeper conceptual connections. They recalled that they began the lesson with limited or surface level knowledge about enzymes and metabolism, often associating metabolism with only food breakdown or understanding enzymes only as general catalysts.

After the lesson, they described understanding how enzyme activity regulates metabolism and varies across individuals to affect medication metabolism. For example, students recalled the following:

> *“This class gave me a deeper understanding of what metabolism and enzymes actually do on a biological level. I used to only know metabolism as the process of breaking down food, but now I understand how the process is actually happening.”*
>
> *“I had a general understanding of enzymes from previous classes, but this unit helped me understand how much the enzyme activity can change…I now understand that the body has so many ways of regulating these enzymes.”*
>
> *“Most biology classes cover this material with little mention of enzymes and metabolism. Most of us knew of these terms, but not how they connected to what we were learning. For example, metabolism was a term we all knew and understood about how the speed of it can affect the body, but not how metabolism and enzymes can affect things like medication.”*
>
> *“Prior to this unit I knew very little about enzymes and metabolism. I knew that enzymes speed up reactions in our body, but I never connected that to them regulating our metabolism.”*
>
> *“I had a very rudimentary knowledge of enzymes prior to this unit, I knew they catalyzed chemical reactions in the body but did not understand that different people have different levels of efficiency for a particular chemical enzyme which can completely change how they react to medicine.”*
>
> *“I knew before that people tended to gain weight when they went on antidepressants but wasn’t really sure why or if it was even a legitimate cause. Learning more about rates of metabolism, and that different rates of metabolism directly affect whether or not people tend to gain weight when using antidepressants solidified for me the link between antidepressants and possible weight gain.”*

#### Perceived utility-value of the lesson

To explore how students perceived the utility of this lesson beyond the classroom, we collected open-ended responses to a question asking how they could apply what they learned to other classes or areas of their life. Utility-value interventions that prompt students to reflect on how aspects of the coursework are useful for their current or future goals can increase students’ motivational beliefs, achievement, and future course-taking intentions^43–45^. Students’ responses indicated that they considered how the material in the lesson was useful for informing their understanding in other courses and how it was useful for helping them make decisions in life.

Other students described the direct personal relevance of how enzymes and metabolism could influence health decisions and everyday lifestyle choices. For example, students described how they viewed the material as being directly relevant for understanding how medications work, anticipating individual differences in medication effectiveness, making informed nutritional and fitness choices, and managing their overall health. Overall, responses suggested that students saw the content as applicable to multiple aspects of their lives. Student responses included:

> *“The fast, slow, and normal metabolizers affect Zoloft gave us a real encounter with how metabolism impacts the effectiveness of different antidepressants. With this new knowledge, we can know why some doses are more or less effective to different people.”*
>
> *“I am interested in pursuing a career in psychology so the metabolism of antidepressants were also interesting to learn about.”*
>
> *“This knowledge helps with understanding how your body processes any medications you may take and how differences in enzymes can affect how drugs are metabolized.”*
>
> *“This knowledge will help me better understand how my body, as well as others’, processes different foods and medications. Understanding enzymes and metabolism helps to clarify why a particular medication may work for me but not for someone else.”*
>
> *“Knowing these things is useful in everyday life because it’s important to understand what you are putting into your body and….knowing how your body responds to drugs.”*
>
> *“I already knew a good bit about enzymes, but I had never connected them to metabolism specifically before. It makes a lot of sense that the enzymes you have such a large impact on your body’s ability to break down different substances makes sense. As someone who takes Escitalopram every day, this has been really helpful for understanding how my own medication works and why it can sometimes take several days before I see side effects if I forget to take it.”*

### Potential Additions or Adaptations to the Lesson

We have implemented this lesson over several semesters and made adjustments based on student feedback and our own observations as instructors. We have refined the activities and the content iteratively to address the areas that students found confusing and/or challenging.

First, we revised the order of topics in the lesson to make it easier for students to follow. In earlier iterations, students learned about SSRI breakdown in the liver before understanding how SSRIs work in the brain. Students found this presentation order to be potentially confusing. We now start an explanation of how SSRIs affect serotonin signaling in the brain, then move into enzyme inhibition, and finish with SSRI metabolism in the liver. This sequence intentionally builds on students’ prior understanding of medication movement from ingestion to elimination and helps content flow more logically on the second day.

Second, we observed that students were also confused by the depiction of Fluoxetine’s (Prozac) function in the slides. Some students felt that the slides made it appear that Fluoxetine inhibited tryptophan hydroxylase, which is not correct. We revised the content on the slides to clarify that tryptophan is the precursor to serotonin and that Fluoxetine acts on the serotonin transporter in the presynaptic neuron to prevent serotonin reuptake, rather than Fluoxetine inhibiting tryptophan hydroxylase. Instructors could also use Omeprazole as a main example for enzyme inhibition. Omeprazole also interacts with tryptophan hydroxylase and enzymes that break down serotonin, which makes it another relevant example for illustrating enzyme inhibition.

On the second day of the lesson, students are asked to answer a question comparing a fast and a slow metabolizer of Prozac. Initially, many students connected metabolism to serotonin reuptake instead of to liver enzyme activity. Reviewing this question as a class helped them make the correct connection, and the revised order of topics should continue to strengthen that understanding.

Another issue on Day 2 was that students had difficulty translating primary scientific data into graphs. Many students were unsure how to represent enzyme activity over time for fast and slow metabolizers taking Fluoxetine. Other students had challenges interpreting what those graphs represented about metabolism and SSRI breakdown. Future iterations of prompts asking students to graph scientific data may benefit from explicitly incorporating graphing instruction into the lesson^23^

An additional challenge was using Google Scholar for the group assignment to locate primary scientific literature. Many students were unfamiliar with how to identify reliable sources or how to recognize primary research articles, despite these topics being addressed during class. We have updated the time allocated on Day 2 to spend even more time reviewing how to use Google Scholar effectively, evaluate credible sources, and to recognize the structure of a primary article.

We also made changes to the list of antidepressants students can choose for their final projects. Specifically, we removed Zoloft and Prozac since they were already covered during class sessions. This change encourages students to examine other SSRIs.

### Institutional Review Board (IRB) Approval

This project was reviewed by the University of Georgia’s Institutional Review Board and determined to be exempt (PROJECT00011312).

## ACKNOWLEDGMENTS

We thank the students who completed the assessments and the peer learning assistants who contributed helpful insights about the challenges students encounter in this lesson.

